# Validation of a PNA clamping method for reducing host DNA amplification and increasing eukaryotic diversity in rhizosphere microbiome studies

**DOI:** 10.1101/2020.05.07.082966

**Authors:** Stephen J. Taerum, Blaire Steven, Daniel J. Gage, Lindsay R. Triplett

## Abstract

Protists and microscopic animals are important but poorly understood determinants of plant health. Plant-associated eukaryotes could be surveyed by high-throughput sequencing of 18S ribosomal RNA (rRNA) genes, but the abundance of plant DNA in rhizosphere samples makes 18S rRNA gene amplification with universal primers unfeasible. Here we applied a pipeline to generate peptide nucleic acid (PNA) clamps to suppress the amplification of plant host DNA during 18S rRNA gene library preparation. We designed a PNA clamp, PoacV9_01, specific to the V9 region of the 18S rRNA gene for plants in the family Poaceae. PoacV9_01 suppressed the amplification of five species of grain crops in quantitative PCR reactions. In an 18S rRNA gene sequencing survey of the rhizosphere of maize, PoacV9_01 reduced the relative abundance of plant reads from 65% to 0.6%, while drastically increasing the number and diversity of animal, fungal, and protist reads detected. Thus, PoacV9_01 can be used to facilitate the study of eukaryotes present in grass phytobiomes. In addition, the pipeline developed here can be used to develop PNA clamps that target other plant species.

## Introduction

Phytobiomes are multi-kingdom communities that significantly impact the health, growth and survival of plants (Huang et al. 2014). Plants host diverse communities of bacteria, archaea, fungi, animals, viruses, as well as autotrophic and heterotrophic protists (Leach et al, 2017). However, sequence-based taxonomic exploration of the phytobiome has focused mostly on its bacterial and fungal components, leaving an underexplored realm of microbial “dark matter” consisting of protists, animals, and viruses. The diversity missing from many studies limits our understanding of microbial food webs and their effects on plant health.

Nonfungal eukaryotic microbes are abundant in the rhizosphere, the zone of soil in direct contact with plant roots, and an important component of the phytobiome (Mendes et al. 2013; Berendsen et al. 2012). In the rhizosphere, plants actively produce exudates that recruit a diverse community of microorganisms (Compant et al. 2019; Mendes et al. 2013; Berendsen et al. 2012), including eukaryotes. A single gram of soil is estimated to contain up to 1 million protists and 100 nematodes, while each square meter can harbor thousands of microscopic arthropods (Leach et al. 2017). The best-understood of these eukaryotes are economically important plant pests, such as parasitic nematodes and *Phytophthora* species, but commensal eukaryotes are also highly influential to ecosystem structure and plant health. Heterotrophic protists and nematodes may shape rhizosphere communities through selective or generalist predation on bacteria and fungi (Jousset et al. 2009; Geisen et al. 2015, 2016), mineralizing nutrients and enhancing the survival and dispersal of beneficial microbes in the process (Zhilei Gao et al. 2019; Rubinstein et al. 2015). Bacterivorous eukaryotes serve as influential hubs in soil microbial networks (Jiang et al. 2017; Xiong et al. 2017), affecting plant biomass, nutritional status, and disease outcomes (Trap et al. 2016; Xiong et al. 2020). Predation may also enhance the beneficial activities of mycorrhizae and plant growth promoting rhizobacteria (Weidner et al. 2017; Henkes et al. 2018), or directly inhibit pathogen growth (Long et al. 2018). Soil arthropods are less-studied in the phytobiome, but are thought to play critical roles in soil nutrient cycling (Parmelee 1995; Lavelle et al. 2006).

High-throughput amplicon sequencing of 16S or 18S rRNA genes is the most widely adopted approach for measuring the bacterial and fungal diversity of the phytobiome. However, there are very few studies profiling non-fungal eukaryotic communities that live in direct contact with plants, such as the rhizosphere. Several factors prevent the broader inclusion of protist and animal taxa in amplicon sequencing studies. For one, many protists and microscopic animals are still uncharacterized on a taxonomic level, although availability of reference sequences for these taxa has increased greatly in recent years. Another complication is that protists are comprised of many high order clades across the eukaryote Tree of Life (Burki et al. 2020; Adl et al. 2019), encompassing far greater taxonomic diversity than fungi or plants alone. Such diversity hampers the ability to design “universal” primers that will amplify all protists and microscopic animals. Group-specific primers have been useful for targeted studies of major protist taxa associated with plants (Ploch et al. 2016, reviewed in Geisen et al. 2019), but these are generally only capable of amplifying a fraction of the eukaryotic portion of the phytobiome.

Several sets of broad-spectrum or universal barcoding primers have been applied to profile diverse protist communities in soil, including those in heavily vegetated and agricultural soils (Mahé et al. 2017; Geisen et al. 2019; Gruyter et al. 2020). Although deep sequencing detects abundant protist taxa, plant DNA interference can greatly limit the sequencing depth of microbial eukaryotes, the number of rare taxa that can be detected, and the number of samples that can be analyzed simultaneously. Root-attached soils contain copious plant DNA from root hairs, border cells, and secreted mucilage (Geisen et al. 2019). In this regard, plant-derived amplification products are likely to dominate the sequencing results. As a result, profiles of the total eukaryotic diversity in direct contact with plants are rare.

One strategy to limit host plant sequence recovery is to block amplification of host DNA using peptide nucleic acid (PNA) clamps, or synthetic oligonucleotides that bind to host targets in a thermally stable manner to block DNA elongation (Karkare and Bhatnagar 2006). This approach has been successfully applied to suppress plastid and mitochondria amplification in studies of the plant bacterial microbiome (Lundberg et al. 2013; Steven et al. 2018), although specialized PNA clamps are needed for some plant species with plastids that differ in the target site (Fitzpatrick et al. 2018). PNA clamps have also been applied to block amplification of host 18S rRNA gene sequences from animals, facilitating surveys of their eukaryotic prey and symbionts (Belda et al. 2017; Terahara et al. 2011). This suggests that amplification of plant host 18S rRNA gene products could also be blocked by PNA clamping. Recently, a study reported the use of a plant-suppressing PNA clamp during sequencing of soil eukaryotes, but no studies on its efficacy, breadth, or potential bias were reported (Gruyter et al. 2020). The true utility of a PNA clamping approach will depend on the degree of host interference that must be overcome, and whether broad-spectrum plant blocking could be achieved without introducing sequence bias.

In this study, we evaluated whether a plant host-suppressing PNA clamp could facilitate surveys of the eukaryotic communities in the phytobiome. A comparative pipeline was applied to identify plant-specific target sequences within universal 18S rRNA gene amplicons, and to design PNA clamps that targeted multiple plant species. A PNA clamp specific to the family Poaceae was selected for validation, and it suppressed amplification of the 18S rRNA gene in maize, rice, wheat, barley, and sorghum. In a sequencing study of rhizosphere soil samples from field-grown maize, addition of the PNA clamp drastically reduced the presence of plant reads and substantially increased the measured number and diversity of non-host eukaryotic sequence variants, including those of fungi, heterotrophic protists, algae, nematodes, and arthropods. The PNA clamp did not introduce any measurable compositional bias. This study demonstrates that the use of a PNA clamp is a highly promising approach for characterizing the eukaryotic phytobiome.

## Materials and Methods

### PNA clamp development

PNA clamps targeting the V4 and V9 hypervariable regions of the 18S rRNA gene were designed following a protocol modified from Belda et al. (2017) and Lundberg et al. (2013). A 1801 bp sequence of the 18S rRNA gene from maize (*Zea mays*) was downloaded from the online SILVA rRNA gene database (accession number LPUQ01000139; Quast et al. 2013). The V4 and V9 regions were isolated after alignment in MEGA v. 7.0.26 (Kumar et al. 2016).

Target regions were fragmented *in silico* into 15, 16 and 17 bp k-mers using the splitter application in EMBOSS (Rice et al. 2000), which was implemented in Jemboss (Carver and Bleasby 2003). Fragments were mapped onto an index of non-plant eukaryote 18S rRNA gene sequences available on the SILVA SSU r132 database using Bowtie v. 1.2.3 (Langmead et al. 2009). The number of times a k-mer matched a sequence in the index was recorded, allowing up to one mismatch in the alignments. Any k-mers that aligned with one or more sequences from the index were removed from consideration.

Candidate k-mers were screened using the online PNA TOOL (https://www.pnabio.com/support/PNA_Tool.htm). Clamps were designed to consist of fewer than 35% guanines and fewer than 50% purines, and a Tm between 76 and 82°C. K-mers that passed screening were mapped to an index of plant sequences from the SILVA SSU r132 database using Bowtie. Two clamp sequences (k-mers) were selected for further testing (see Results): PoacV4_01 (V4 region, sequence 5’-TCGGTTCTCGCCGTGA-3’) and PoacV9_01 (V9 region, sequence 5’-GCCGCCCCCGACGTC-3’). The PNA clamps were synthesized at PNA Bio, Inc. (Newbury Park, CA), and resuspended in nuclease-free water to a concentration of 100 µM.

### Plant lines, protist isolates, and growth conditions

Seeds of maize B73, rice (*Oryza sativa*) *L.* cv. Kitaake, wheat (*Triticum aestivum*) *L.* cv. Byrd, barley (*Hordeum vulgare*) *L.* cv. Morex, and sorghum (*Sorghum bicolor*) *L.* cv. BT623, *Arabidopsis thaliana* ecotype Col-0, and *Nicotiana benthamiana* were germinated in wetted Turface (Turface Athletics, Buffalo Grove, Illinois, USA) in a greenhouse (28°, 14h days). After one week, the stems and leaves of the seedlings were collected.

Protist DNA was isolated from two laboratory lines (Daniel Gage Lab, University of Connecticut) isolated from soybean rhizospheres: UC1 (*Colpoda* sp.) was isolated in Mansfield, Connecticut, in 2013, while UC5 (*Cercomonas* sp.) was isolated in Columbia, Connecticut, in 2015. Both cultures were initiated from single cells into soil extract medium (20 mg KH_2_PO_4_, 20 mg MgSO_4_•7H_2_O, 200 mg KNO_3_, 100 mL soil extract per liter, Culture Collection of Algae and Protozoa, Dunstaffnage Marine Laboratory, UK). Cultures were maintained using heat-killed *E. coli* (DH5a, OD_600_ = 0.005) as a food source.

### Initial PCR tests of clamp function

DNA was extracted from plant tissue using the GeneJET Plant Genomic DNA Purification kit (Thermo Scientific, Waltham, Massachusetts, USA). Protist DNA was extracted using the GeneJET Genomic DNA Purification kit (Thermo Scientific). DNA concentrations were measured using the Qubit dsDNA HS kit (Thermo Scientific), after which the samples were each diluted to 1 ng/µL. An initial set of PCR reactions was conducted on the plant and UC1 templates. The V4 region was amplified using the primers TAReuk454FWD1 (5’-CCAGCASCYGCGGTAATTCC-3’) and TAReukREV3 (5’-ACTTTCGTTCTTGATYRA-3’; Stoeck et al. 2010), while the V9 region was amplified with the Earth Microbiome Project primers Euk1391F (5’-GTACACACCGCCCGTC-3’; Lane 1991) and EukBr (5’-TGATCCTTCTGCAGGTTCACCTAC-3’; Medlin et al. 1988). Reactions consisted of 1 unit Invitrogen Taq DNA polymerase (Thermo Scientific), 1x PCR buffer, 0.5 µM forward and reverse primers, 0.2 mM dNTPs, 1.5 mM MgCl2, and 0.2 ng/µL of template in a 25 uL reaction volume. Clamps PoacV4_01 or PoacV9_01 were added to reach the following concentrations: 0, 1.5, 7.5, and 15 µM. Cycling conditions were 3 minutes at 94°C, followed by 30 cycles of 94°C for 45 seconds, 55°C for 30 seconds, and 72°C for 90 seconds, followed by a final extension of 72°C for 10 minutes. The PCR amplicons were separated on a 1% agarose gel and visualized after staining with ethidium bromide.

### qPCR test of PNA clamps

qPCR reactions of the V4 and V9 regions were conducted using the SsoAdvanced Universal SYBR Green Supermix (Bio Rad, Hercules, California, USA), and the same primers used for PCR reactions. Reactions consisted of 1x Supermix, 0.5 µM forward and reverse primers, and 0.2 ng/µL of template in a 10 uL reaction volume. For initial qPCR tests, the PNA clamps were added to a final concentration of 1.5 µM. Cycling conditions for amplifying V4 were 2 minutes at 95°C, followed by 40 cycles of 95°C for 10 seconds and 60°C for 60 seconds, while cycling conditions for V9 were 2 minutes at 95°C, followed by 40 cycles of 95°C for 10 seconds and 60°C for 15 seconds. Reactions were conducted on a CFX96 Touch Real-Time PCR machine (Bio Rad).

Further studies focused on optimizing and testing the V9 clamp, PoacV9_01. To determine the optimal concentration of this clamp, qPCR reactions were conducted as described above, except that PoacV9_01 was added to reach the following concentrations: 0, 0.75, 1.5, 3.75, and 7.5 µM. Each PNA concentration was tested on maize DNA in triplicate. 3.75 µM was found to be the optimal concentration of PoacV9_01, and this concentration was used in all subsequent experiments.

To determine the efficacy of PoacV9_01 in blocking predicted target and nontarget organisms, qPCR reactions were then conducted as described above using the DNA isolated from a selection of plants and protists. Water was used as the negative control. Reactions were conducted in triplicate using a PNA clamp concentration of 3.75 µM. Cycle threshold (C_T_) values were statistically compared between clamp and no clamp reactions using a Student’s t-test.

### Preparation and high throughput sequencing of maize rhizosphere libraries

We next studied the impact of the clamp on high-throughput sequencing of the maize rhizosphere. B73 maize was planted on May 15, 2019, in Griswold Farm in Griswold, Connecticut, and May 17, 2019, in Lockwood Farm in Hamden, Connecticut. Root crowns of four individual plants, or two from each farm, were harvested at 8 weeks after planting (V6 stage) using a drain spade. Root crowns were shaken vigorously for 1 minute, and the top 10 cm from 5 assorted roots were collected using sterile scissors and forceps and placed on ice. Rhizosphere soil samples were collected within 30 minutes of root collection by vortexing 2 minutes in 35 mL phosphate buffered saline following the protocol of McPherson et al. (2018) and stored at −80°C until DNA was extracted. The two samples collected at the Hamden, CT site were labeled L1 and L2, and the samples from Griswold site were G1 and G2.

Rhizosphere DNA was extracted using the DNeasy PowerSoil Kit (Qiagen, Germantown, Maryland, USA). Two separate V9 amplicon libraries were generated for each sample: one with the PNA clamp, and one without. Preparation of the eight libraries for high-throughput sequencing involved two separate PCR steps using Invitrogen Platinum *Pfx* DNA polymerase (Thermo Fisher). In the first set of reactions, PCR amplification was conducted using Euk1391F and EukBr primers with the Illumina sequencing adaptors (Euk1391F: TCGTCGGCAGCGTCAGATGTGTATAAGAGACAGGTACACACCGCCCGTC; EukBr: GTCTCGTGGGCTCGGAGATGTGTATAAGAGACAGTGATCCTTCTGCAGGTTCACCTA C) at a final concentration of 0.5 µM. Clamp reactions were conducted with the PNA clamp at a final concentration of 3.75 µM. The thermal cycler protocol consisted of a denaturation step of 3 minutes at 95°C, followed by 25 cycles of a denaturation step of 30 seconds at 95°C, an annealing step of 30 seconds at 55°C, and an extension step of 30 seconds at 72°C, followed by a final extension step of 5 minutes at 72°C. After amplification, the libraries were cleaned using the GeneJET Gel Extraction and DNA Cleanup Micro kit (Thermo Scientific). In the second set of reactions, the samples were barcoded on both ends using Nextera DNA CD Indexes (Illumina Inc., San Diego, California, USA). The thermal cycler protocol consisted of a denaturation step of 3 minutes at 95°C, followed by 8 cycles of a denaturation step of 30 seconds at 95°C, an annealing step of 30 seconds at 55°C, and an extension step of 30 seconds at 72°C, followed by a final extension step of 5 minutes at 72°C. The barcoded libraries were cleaned one final time using the GeneJET Gel Extraction and DNA Cleanup Micro kit.

The eight libraries were quantified using the Qubit dsDNA HS kit, and then diluted to the same concentration. The libraries were pooled and sequenced on the Illumina iSeq 100 platform. The samples were sequenced using 2 x 150 bp chemistry. The raw sequence data have been submitted to the NCBI Short Read Archive database under accession number NCBI: PRJNA630266.

### Bioinformatic analyses

#### Quality control and filtering

Read filtering and assembly were conducted using the mothur v. 1.44.0 package (Schloss et al. 2009). Paired-end demultiplexed sequences were assembled into contigs. Contigs were screened to retain sequences between 100 and 200 bp in length, with no ambiguous bases, and a maximum homopolymer length of eight. Filtered contigs were aligned to the SILVA v. 132 SSU reference library. Chimeric sequences were identified using VSEARCH (Rognes et al. 2016) as implemented in mothur and all potentially chimeric sequences were removed.

#### Taxonomic classification

Sequences were assigned to Amplicon sequence variants (ASVs, sequences sharing 100% sequence similarity). We examined ASVs instead of operational taxonomic units (OTUs), because ASV methods have a potentially higher sensitivity and resolution (Callahan et al. 2017), which would allow us to better assess the impact of PoacV9_01 on the detection of taxonomic diversity. Representative ASVs were classified using the naïve Bayesian classifier in mothur (Wang et al. 2007) against the SILVA v. 132 SSU reference database (Quast et al. 2013). ASVs were also separately classified against the Protist Ribosomal Reference (PR^2^) database (Guillou et al. 2013). Classifications with a bootstrap of 80% or higher were retained.

#### Analysis of abundance and diversity

ASV abundance data were merged with the SILVA and PR^2^ taxonomic classification files using the phyloseq package (McMurdie and Holmes 2013) implemented in R. The SILVA and PR^2^ ASV classifications were used to compare ASV abundance between corresponding clamped and unclamped libraries. Rank abundance curves were generated by sorting non-plant ASVs by abundance in R and plotting in ggplot2. Alpha-diversity statistics were calculated using mothur. Rarefaction curves of the ASVs in each library were generated using the vegan package (Oksanen et al. 2019) in R.

## Results

### Implementation of a pipeline for design of PNA clamps

To facilitate sequencing of the eukaryotic rhizosphere microbiome, we sought to design a PNA clamp that would bind specifically to plant sequences, but not non-plant sequences. We targeted two commonly used 18S rRNA gene universal taxonomic primer sets targeting the V4 and V9 hypervariable regions, respectively. To design such a clamp, we implemented a k-mer comparative alignment pipeline adapted from previous work (Fig. 1; Lundberg et al. 2013).

**Fig. 1.**
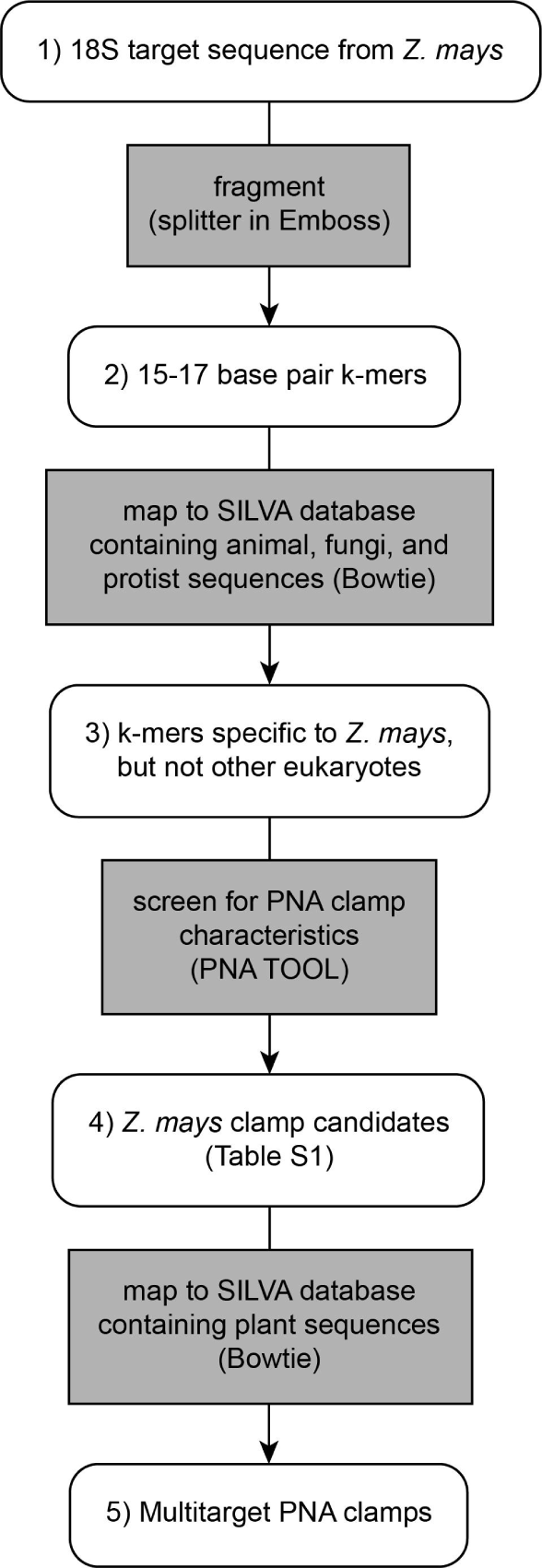
Flowchart depicting the computational pipeline utilized in this study, which can be adapted for future PNA clamp design. Gray square boxes denote processes, rounded white boxes indicate inputs and outputs.

#### Selection and *in silico* analysis of clamps

Nineteen V4 k-mers and one V9 k-mer met the design criteria for effective PNA clamps (Table S1). No maize-specific k-mers from either region mapped to all or most plant sequences in the index, but three V4 k-mers mapped to three species in the family Poaceae and one species in the family Asparagaceae, while the one V9 k-mer mapped to 22 species in the family Poaceae, one species in the family Asparagaceae, and one species in the family Xyridaceae. One k-mer from each variable region was selected as a PNA clamp sequence (Fig. S1); these were named PoacV4_01 (V4 region, sequence 5’-TCGGTTCTCGCCGTGA-3’) and PoacV9_01 (V9 region, sequence 5’-GCCGCCCCCGACGTC-3’). These were selected because they were predicted *in silico* to bind multiple plant species (Table S2). PoacV9_01 had a 100% match with several other model grain species (Fig. 2), and only differed by one to two bases from the target region of some non-grass plants that were examined, suggesting small modifications to the sequence could make it applicable for studies in other host plants.

**Fig. 2.**
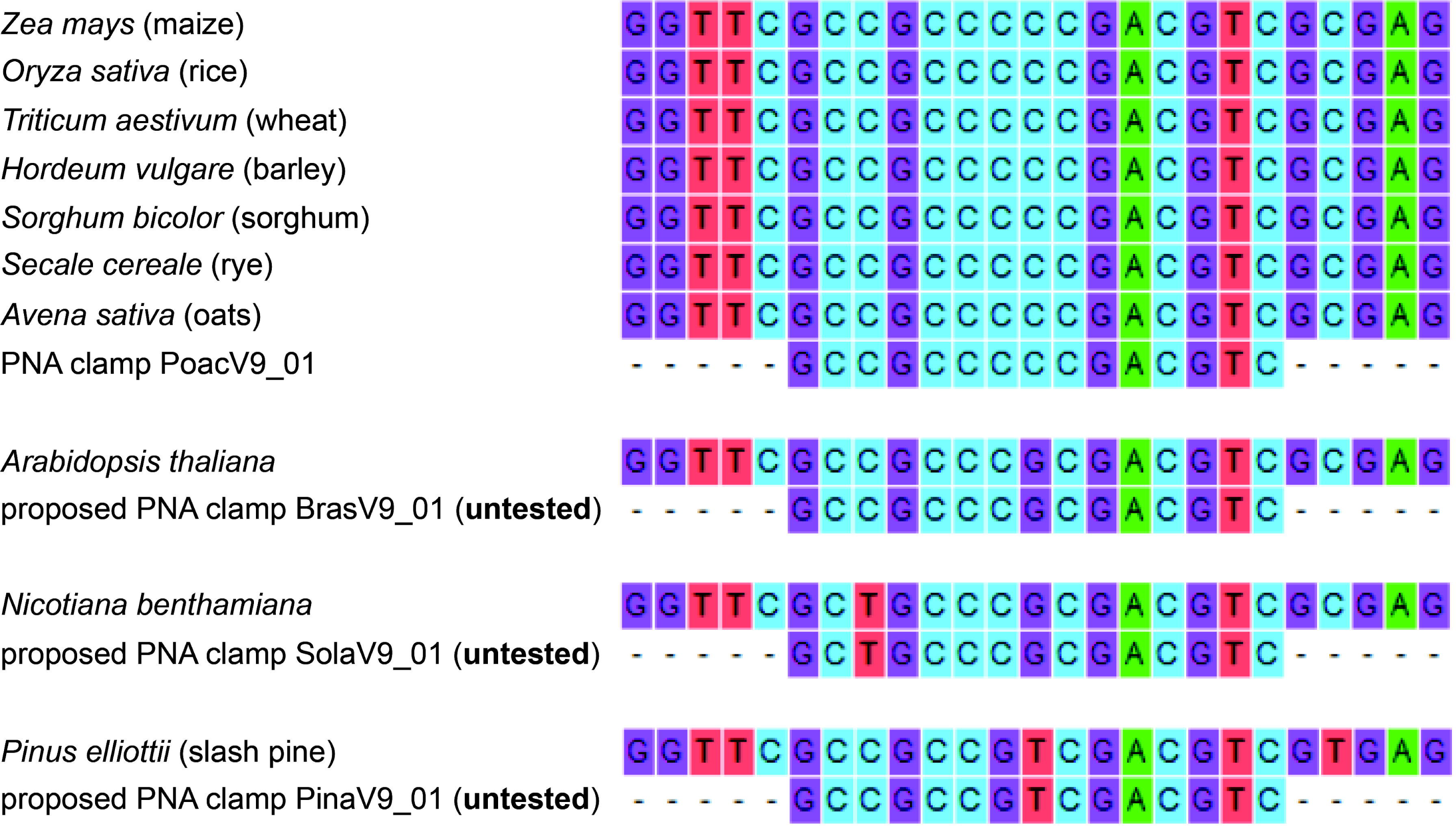
18S rRNA gene fragments of select plant species that align with the PNA clamp developed in this study (PoacV9_01). The clamp was experimentally demonstrated to effectively block the amplification of DNA from maize, rice, wheat, barley, rye and sorghum, and should be effective for other grass species, such as oats. Small modifications in the clamp sequence should make it effective for other plant species, such as *Arabidopsis thaliana, Nicotiana benthamiana*, and *Pinus elliottii*. The proposed clamps for *A. thaliana, N. benthamiana*, and *P. elliottii* have not been tested on plant material, but have been tested *in silico* following the protocol outlined in this study (see Materials and Methods).

#### Preliminary qualitative PCR evaluation of clamp function

Initial conventional PCR tests using 18S rRNA gene universal primers in clamp and no-clamp reactions demonstrated that both clamps suppressed amplification of DNA from maize, as seen by a reduced band intensity compared to a no-clamp reaction (Fig. S2). Neither clamp suppressed amplification from the protist UC1. Both clamps also suppressed amplification of rice DNA, and PoacV9_01 appeared to suppress amplification of sorghum and *A. thaliana* as well. Although these results were only visualized at the end of 30 cycles and were not quantitative, they suggested that both PoacV4_01 and PoacV9_01 could effectively block 18S rRNA gene amplification from maize and rice.

### Optimization and validation of the V9 PNA clamp PoacV9_01

To quantify the efficiency of blocking by the PNA clamps, we followed up these initial tests with qPCR reactions. Unfortunately, over numerous attempts we were unable to obtain consistent qPCR amplification of the maize V4 region using the primer set TAReuk454FWD1 and TAReukREV3, which has a longer amplicon than recommended for qPCR using fluorescent staining methods. Because of the broader potential species range of the PoacV9_01 clamp, as well as the advantages of the universal V9 primer set to our planned future studies (see Discussion), we decided to focus on PoacV9_01 for in-depth analyses.

PNA clamp concentration may affect its efficacy (Lundberg et al. 2013; Belda et al. 2017). In a test of several clamp concentrations, PoacV9_01 suppressed the amplification of maize DNA by approximately 5 cycles at concentrations as low as 0.75 µM (Table S3). 3.75 µM increased the suppression by approximately 1 more cycle, and with little to no increased efficiency at higher concentrations (Table S3). Therefore, 3.75 µM was selected as the optimal PNA concentration for further analyses.

Next, we tested the efficacy of the PoacV9_01 clamp in suppressing amplification from five *Poaceae* crops that have an identical match to the V9 target sequence: maize, rice, wheat, barley, and sorghum. In addition, we evaluated suppression of the model plant *A. thaliana*, predicted to have only one mismatch in the eighth nucleotide of the target sequence (Fig. 2), and *N. benthamiana*, predicted to have two mismatches in the third and eighth nucleotides. DNA from protist isolates UC1 and UC5 were also tested to evaluate potential off-target suppression of non-plant eukaryotes. PoacV9_01 was highly effective in suppressing the amplification of all five Poaceae crops (Table 1). Clamp addition delayed fluorescence detection in these five species by 5.35 to 9.12 cycles compared to the corresponding unclamped reactions. There was slight suppression of amplification of the *A. thaliana* V9 region, although much less efficient than that observed with the grass species (Table 1). In contrast, PoacV9_01 had no significant effect on amplification of protist or *N. benthamiana* DNA. These results indicate that the PNA clamp PoacV9_01 effectively suppresses amplification from grass species with matching V9 target sequences, and may reduce amplification from species with a single central mismatch, but does not suppress amplification from species with two or more mismatches.

**Table 1.**
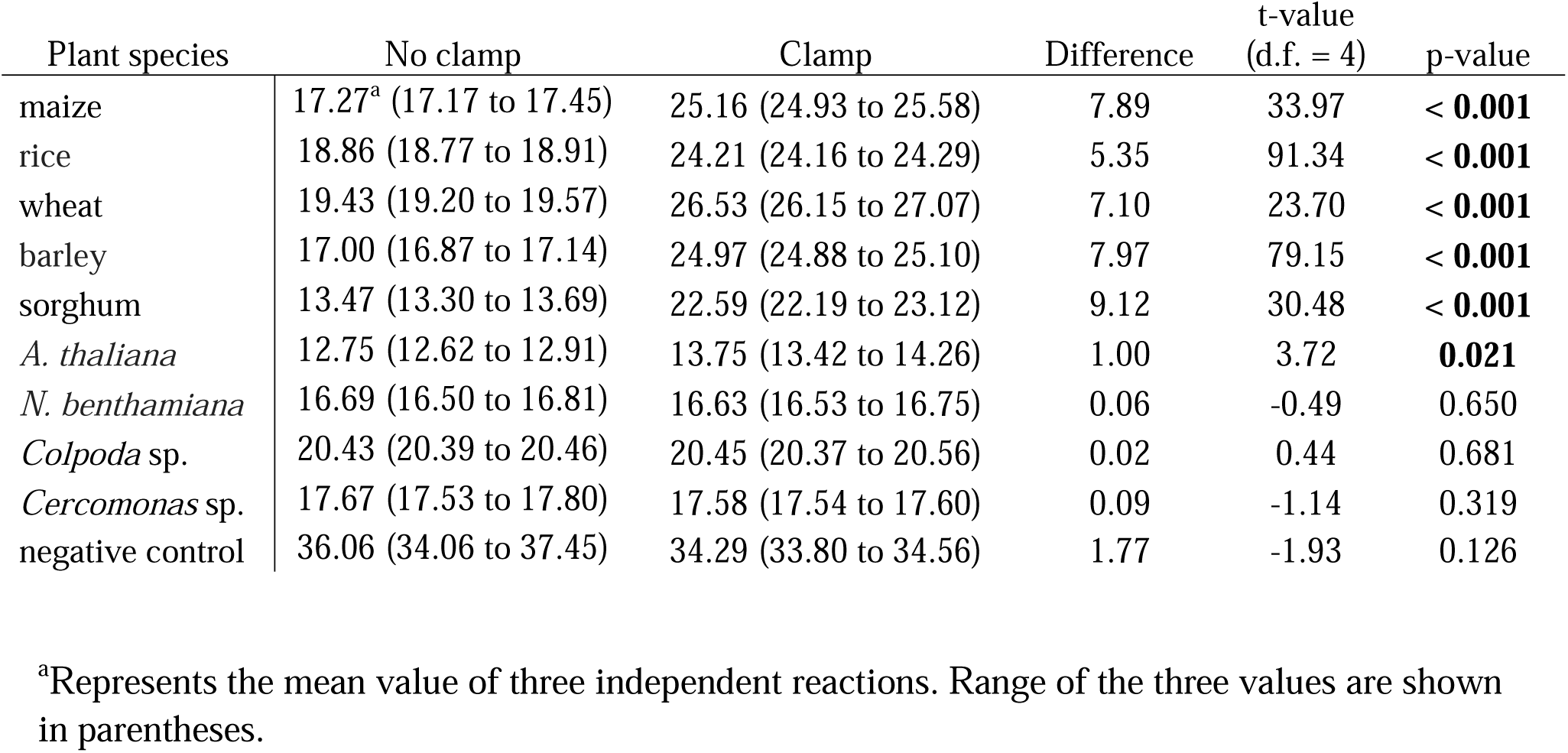
Mean cycle thresholds and ranges observed in 18S V9 amplification of selected species, with and without addition of the PNA clamp Poac9V_01 at 3.75 µM.

### Clamp PoacV9_01 reduces amplification of host DNA and increases ASV abundance and diversity

Although PoacV9_01 suppressed amplification of purified maize DNA in a qPCR assay, it was still uncertain how effective the clamp would be in the context of sequencing a complex rhizosphere sample, or whether clamp addition would introduce a bias to amplification and recovery of sequences. To answer these questions, we applied the PoacV9_01 clamp to high-throughput sequencing of rhizosphere soil samples collected from four field-grown maize plants. Clamp and no-clamp libraries were simultaneously prepared and sequenced for each plant, in order to determine the effect of PoacV9_01 on sequence recovery within each sample.

After quality filtering we recovered 2,291,527 sequences (286,441 average per sample). These sequences could be clustered into an average of 44,286 ASVs per sample.

### PoacV9_01 drastically reduced the sequencing of host reads in rhizosphere samples

Representative ASV sequences were classified according to two taxonomic databases commonly used in eukaryote profiling studies, SILVA (Quast et al. 2013) and PR^2^ (Guillou et al. 2013). The PR^2^ database includes a more comprehensive catalog of eukaryotic taxa, especially protists (Guillou et al. 2013), so we hypothesized that this database would allow us to classify a higher proportion of eukaryotic sequence variants or produce classifications to deeper taxonomic ranks. In this manner, we could also assess if the classification database employed would affect the conclusions from a sequence-based study of eukaryotic diversity.

Regardless of the classification database, addition of PoacV9_01 reduced the mean proportion of plant-derived reads from 66.5% to 0.6% (Fig. 3A). The sequencing recovery of other eukaryotes was increased correspondingly (Table S4), yielding a much higher proportion of animal, fungal, and protist reads in the clamped libraries than the unclamped libraries. The largest difference between reads classified by the two databases was in the bacteria and archaea (Fig 3A). As these organisms are not included in the PR^2^ database they are presumably represented the bin “unclassified” (Fig 3A). Thus, these data suggest there is some amplification of bacteria and archaea from the rhizosphere samples.

**Fig. 3.**
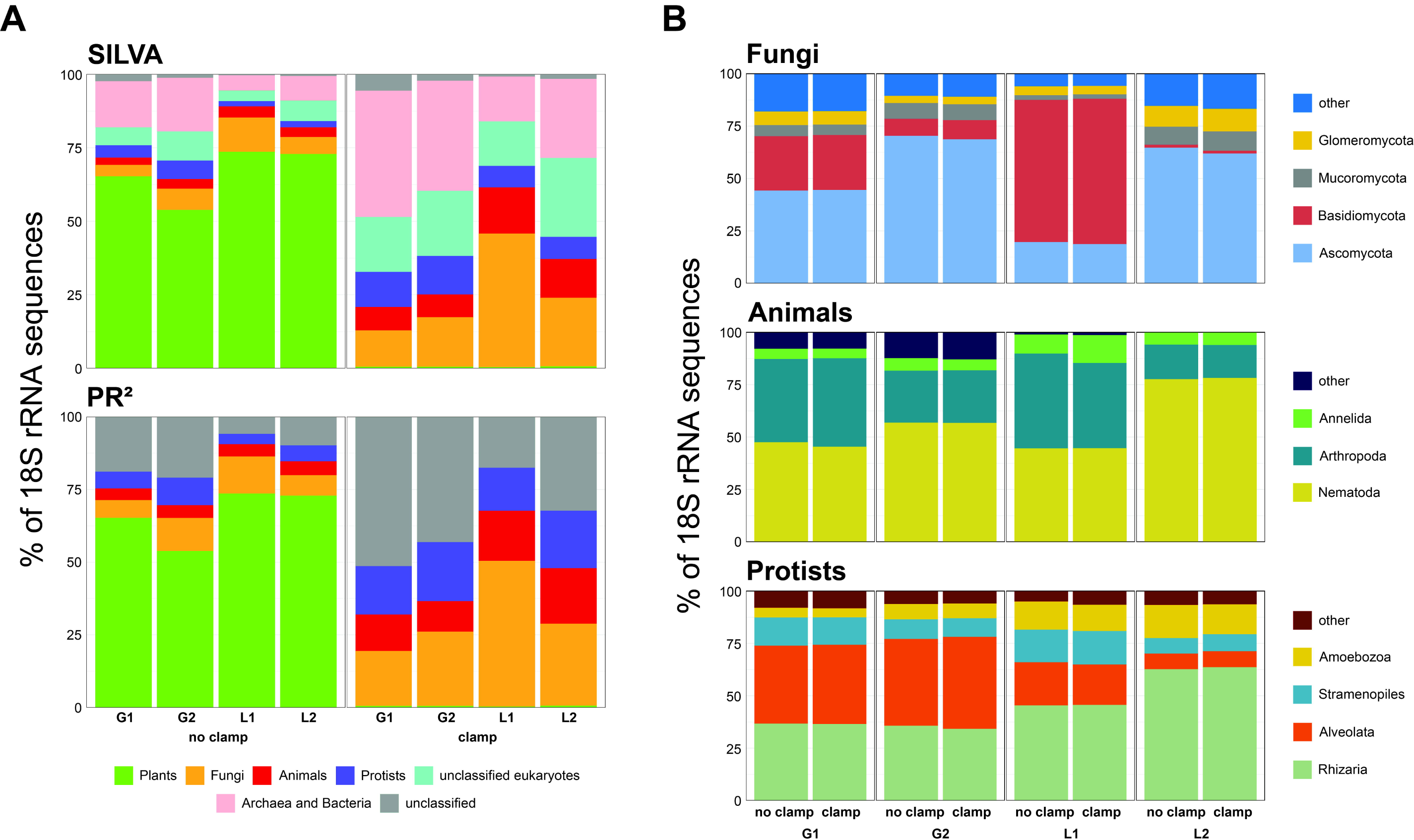
Bar charts showing the relative abundance of taxonomic bins in the sequence datasets. **A.** The relative abundances of kingdom level groups found in each rhizosphere sample, using either the SILVA or PR^2^ database for classification. The bars, each of which represent a sequenced library, are organized by clamp treatment. B. The relative abundances of kingdom level groups of non-plant eukaryotes found in this study, as identified using the PR^2^ database. The bars are organized by sample I.D. The category “other” is the sum total of the taxonomic groups not displayed.

A higher proportion of sequences were classified as animals (14.9%), fungi (30.7%) and protists (17.9%) by the PR^2^ database than by the SILVA database (11.1% to the animals, 24.5% to the fungi, and 10.0% to the protists) suggesting that PR^2^ may be more discriminating in the classification of eukaryotes in the phytobiome. Despite these differences, the patterns between datasets were maintained. For example, both databases identified that the relative abundance of fungi was highest in sample L1. Taken together, these data suggest that the differences between classification databases are relatively small, but potential biases in how the databases assign taxa should be taken into consideration before selecting one database over the other (Dupont et al. 2016).

Using the PR^2^ database, which classified more sequences to Eukaryotic groups than SILVA, we examined the major non-plant groups at deeper taxonomic levels. There were only very minor differences in the proportional abundances of animal, fungal and protist sequences between the clamp and no-clamp libraries (Fig. 3B, Table S5), suggesting that PoacV9_01 is not a source of bias. However, there were large differences among the four samples. Ascomycetes were the dominant fungal group within samples G1, G2 and L2, while basidiomycetes were the dominant fungal group within sample L1. In addition, nematodes were the dominant animal group in samples G2 and L2, while they were codominant with arthropods in samples G1 and L1. Finally, alveolates were the dominant protist group in samples G1 and G2, while rhizarians were the dominant protist group in L1 and L2. Some of these patterns could be the result of differences in biogeography or soil chemistry (e.g., the differences in protist abundance between the Griswold and Hamden samples). In general, though, the four plants harbored very different communities of eukaryotic microbes. Studies with more replicates and deeper sequencing are needed to detect and verify any ecological patterns.

### PoacV9_01 increased the abundance and diversity of 18S rRNA gene sequences recovered from rhizosphere soils

Next, we asked whether addition of the clamp increased the number and diversity of taxa detected by sequencing. ASV-based analysis demonstrated that PoacV9_01 increased the number of ASVs in each library by 15,161 to 38,330 compared with the corresponding unclamped library (Table 2), or an increase of 50-100%. Across samples, we detected a mean of 8,832 additional ASVs from fungi, 7,432 from protists, and 4,604 from animals that were not detected in the absence of PoacV9-01 (Table S6). The increases in detected diversity were reflected in the Shannon diversity indices for each sample, which were doubled in the clamp libraries relative to the no-clamp libraries (Table 2). Good’s coverage estimator, an estimate of the proportion of total species represented in a sample, was consistently lower in the clamp libraries than in the no-clamp libraries, suggesting that deeper sequencing of the clamp libraries would result in the detection of many more ASVs compared with the no-clamp libraries. Good’s coverage for the clamped samples ranged from *ca.* 86 to 89%, suggesting that the majority of the expected genetic diversity of the Eukaryotic community was recovered with this sampling depth. The higher number of ASVs detected in the clamp samples was supported by rarefaction curves (Fig. 4), which confirmed that ASV counts were consistently higher for clamp samples than for no-clamp samples across the samples. However, the rarefaction curves did not plateau, even in the clamped samples, suggesting that even with the increased recovery of ASVs with the PNA clamps the total diversity of the eukaryotic community has only been partially sampled.

**Table 2.**
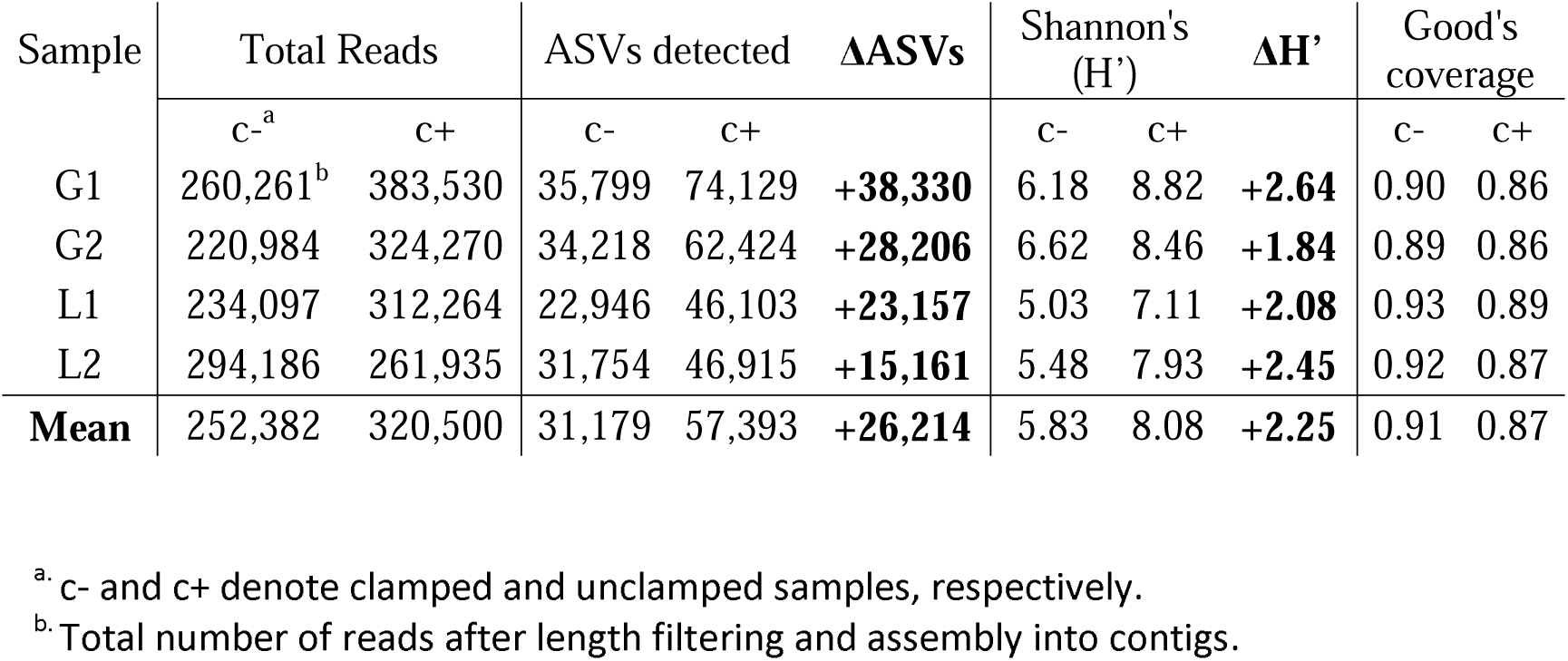
Alpha diversity statistics for the no-clamp and clamp libraries.

**Fig. 4.**
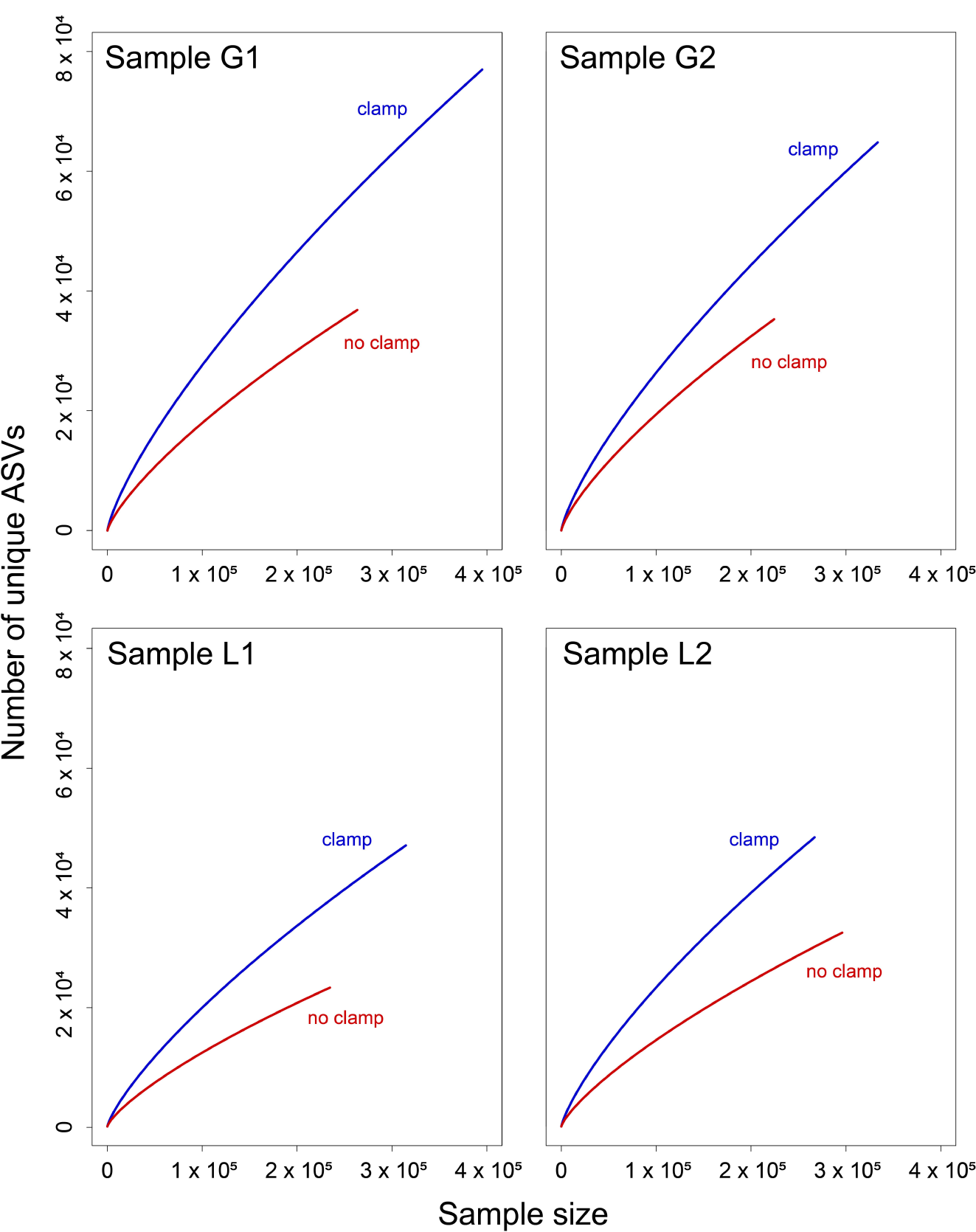
Rarefaction curves of the libraries generated from the four rhizosphere samples. Curves represent the number of ASVs occurring in arbitrary samples of increasing size in clamp and no-clamp libraries for each sample.

### PoacV9-01 did not bias ASV recovery

We next asked whether the PoacV9-01 clamp might introduce bias into sequencing results, potentially through off-target binding, increasing amplification bias, or other mechanisms. After bioinformatically removing plant reads from the libraries, rank abundance curves were plotted for each sample (Fig. 5). As can be seen in Figure 5, ASVs consistently had the same rank abundance relationship between clamped and non-clamped samples. In other words, the most abundant ASVs in the non-clamped samples were consistently identified as the most abundant ASVs in the clamped sample. Furthermore, the ASVs were generally present in similar relative abundances, suggesting that the community profile for the dominant eukaryotic members was unaffected by including a PNA clamp in the PCR amplification.

**Fig. 5.**
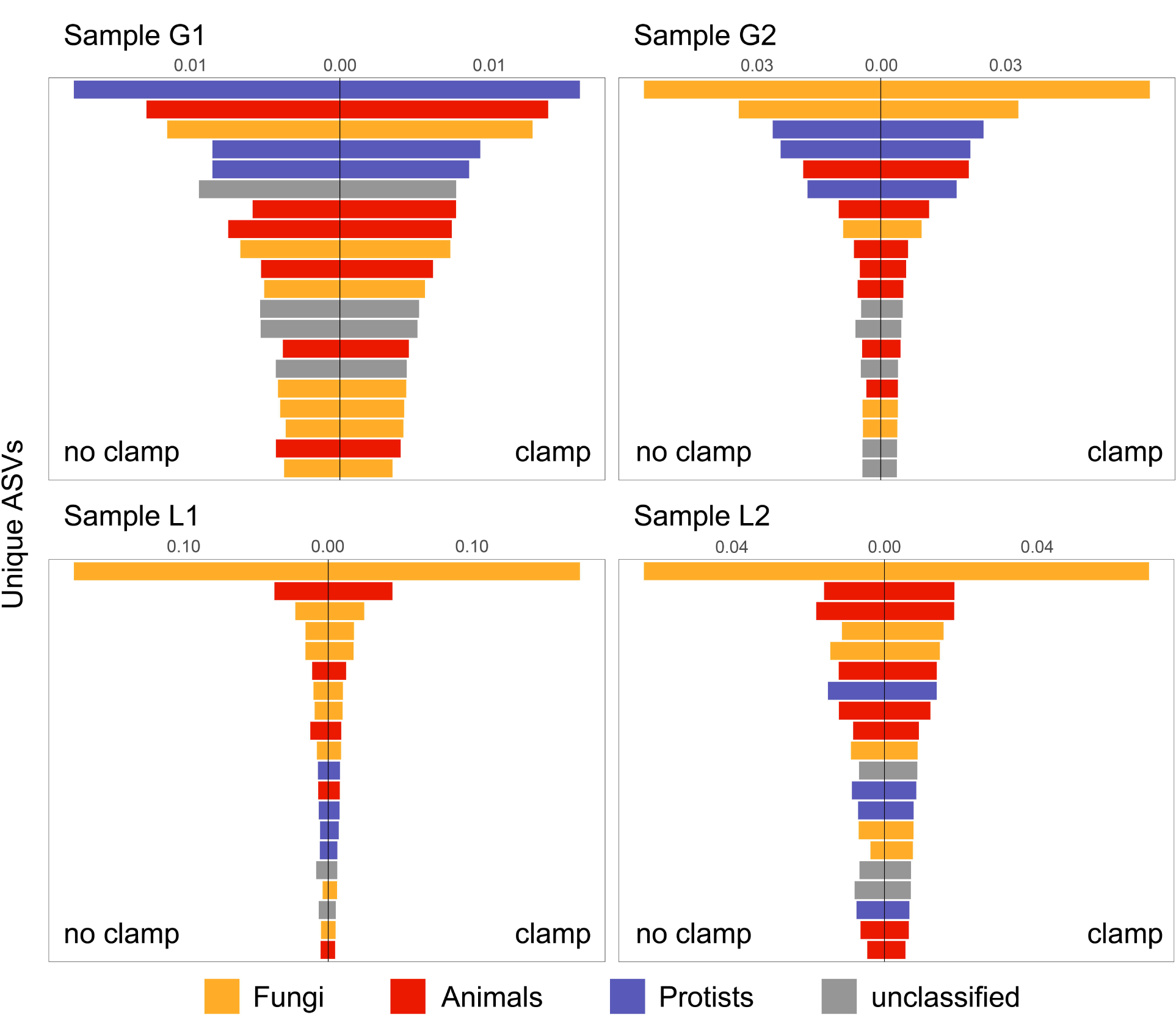
Rank abundance curves of the libraries generated from the four rhizosphere samples, after plant sequences were removed. Each bar represents a single ASV. The 20 most abundant ASVs in each sample are shown in order of their ranking in the clamp library. In this respect, the absence of an ASV in a sample means that it was not among the 20 most abundant ASVs, not absent from the sample. Bar lengths represent ASV relative abundance with the clamp (right side x-axis) and without the clamp (left side x-axis). ASVs are colored according to their kingdom level taxonomic classification based on the PR^2^ database.

An additional observation that arose from this analysis is that the community profile of ASVs between individual plants was largely unique. For instance, the dominant ASV for three of the plants was a fungus, but for one plant a protist was the most abundant ASV (Fig. 5). In fact, only one ASV, identified as a nematode in the genus *Aporcelaimellus*, was conserved amongst the most dominant ASVs in the samples. These data support that the Eukaryotic communities are complex, and deep sequencing with highly replicated sampling will be required to disentangle the ecological relationships between plants and their phytobiomes.

## Discussion

The majority of plant microbiome studies have focused on bacteria and fungi, in part because the abundance of host DNA in plant associated samples limits the sequencing from other eukaryotes. Here we optimized a pipeline (Belda et al. 2017; Lundberg et al. 2013) for the design of PNA clamps to suppress the amplification of plant 18S rRNA gene sequences during high-throughput sequencing. We developed and validated the PNA clamp PoacV9_01, which suppressed the amplification of 18S rRNA genes from five species of model grain crops. In high-throughput sequencing analysis of maize rhizosphere DNA, the clamp eliminated *ca.* 99% of host sequence interference, reducing the proportion of plant reads from 66.5% to 0.6%. Incorporation of PoacV9_01 increased the measured diversity of other eukaryotes including animals, fungi and protists, increasing the number of sequence variants without biasing the composition of eukaryote groups. This study establishes PNA clamping as a promising method to facilitate profiling of eukaryotes in the phytobiome.

Addition of the PoacV9_01 clamp confers several advantages to high-throughput sequencing of plant-associated eukaryotes. By greatly increasing the number of non-plant reads and ASVs detected across the samples, clamp addition substantially increased the value of a relatively small-scale sequencing study. This demonstrates that PoacV9_01 or other PNA clamps could greatly increase the depth and cost-effectiveness of eukaryotic phytobiome studies, allowing researchers to study a greater number of samples simultaneously. PoacV9_01 could also allow researchers to obtain more informative data from relatively inexpensive sequencing platforms, such as the Illumina iSeq 100 platform used in this work. The increase in sequencing efficiency could also make universal 18S rRNA gene primers a viable broad-scope alternative to group-specific primers, including those commonly used in studies of phytobiome fungi (Hannula et al. 2017; Zhiyuan Gao et al. 2019). Studies are underway to determine whether the approach described here compares favorably to fungal-specific primers in measuring community composition.

An initial goal of this study was to identify PNA clamps that would universally block 18S rRNA gene amplification in plants without inhibiting the amplification of other eukaryotes. However, protists are extremely diverse, and our pipeline did not find a universal plant clamp in the V4 or V9 regions that would not also block at least some other eukaryotic taxa. This was expected, as even “universal” plastid-blocking clamps most commonly used in microbiome studies are not universal to all plants (Fitzpatrick et al. 2018). However, *in-silico* analysis showed that the PoacV9-01 clamp is modifiable; that is, correcting the sequence to match the same V9 site in other plant families results in strong candidate clamps for a variety of model species (Fig. 2). Work is underway in our group to validate the efficacy and breadth of these modified clamps in other plant families. While this targeted clamping strategy will limit the use of PNA clamping on mixed plant ecosystems or in weed-infested soils, the approach will still be applicable for the majority of agricultural phytobiome studies performed on monoculture crops. Furthermore, because this study is focused on sequencing rhizosphere samples from an individual plant, the presence of other plant DNA is likely limited. If multiple species suppression is desired, multiplexing could also be a viable approach to suppress the amplification of multiple targets at once; this strategy has been applied to suppress different plastid and mitochondrial targets from plants for bacterial microbiome studies (Lundberg et al. 2013). It’s also possible that, as more becomes known about the eukaryotic phytobiome, it will be considered acceptable to design a clamp that is more universal to plants, but which also suppresses some protist groups of lesser interest.

Despite its targeted nature, PoacV9_01 effectively suppressed qPCR amplification from five major grain crops: maize, rice, wheat, sorghum, and barley. These species represent five genera from three distinct subfamilies in the Poaceae, and the target region was confirmed *in silico* to be conserved in other Poaceae *sp.* (Fig. 1). Therefore, we anticipate that PoacV9_01 will be useful across varieties within these species, as well as with other grasses such as rye or oats. However, we recommend that the clamp should be optimized for each new host species prior to a sequencing run. PoacV9_01 also suppressed the amplification of *A. thaliana* to a small but significant degree. This supports prior findings that PNA clamps can still affect amplification of targets with a single mismatch (Terahara et al. 2011), but also demonstrates that a central mismatch greatly reduces suppression efficiency.

This study sought to design clamps that would work for common universal primers targeting the V4 and V9 hypervariable regions of the maize 18S rRNA gene. Both the V4 and V9 regions are widely used in eukaryotic community studies, although they carry different biases and may result in highly distinct results when applied to the same sample (Giner et al. 2016; Tragin et al. 2018; Hirakata et al. 2019). We ultimately focused on developing a clamp for the V9 region in this work, for three reasons: first, the candidate V9 clamp PoacV9_01 matched a wider predicted species range than any clamp designed for the V4 region, making it more broadly applicable for downstream studies. Second, the V9 region is much more consistent in length than the V4 region, which can range from 230 to up to 520 bp in eukaryotes (Nickrent and Sargent 1991). The V9 length homogeneity may limit amplification bias (Geisen et al. 2018). The shorter length of the V9 amplicon also allows full sequence coverage from paired ends, providing more confidence to ASV methods. Finally, the V9 primers Euk1391F and EukBr primers are a standard tool of the Earth Microbiome Project, a global microbiome cataloguing effort aiming to sequence the microbial diversity of ∼200,000 locations globally (Gilbert et al. 2014), potentially permitting plant microbiomes to be compared with other communities.

The pipeline also produced a proposed V4 clamp, PoacV4_01, predicted to target maize. The V4 amplicon has its own advantages for many applications; for example, its longer size makes it potentially more phylogenetically informative than the V9 region (Geisen et al. 2018). The V4 region is also better represented in the SILVA and PR^2^ databases (Geisen et al. 2018), and has been previously adopted as the barcode for protist taxonomic analysis (Pawlowski et al. 2012). The universal V4 primers used here are also less likely to result in off-target amplification from bacteria, a known issue with Euk1391F and EukBr (Kounosu et al. 2019) that was also evident by the bacterial sequence in our study. PoacV4_01 reduced apparent amplification from rice and maize in conventional PCR, and is a promising candidate for sequencing studies in those species. However, PoacV4_01 has not been validated in a quantitative manner, and we would caution potential users to perform optimization studies before implementing this clamp in a sequencing reaction.

Despite its importance to plant health, the nonfungal eukaryotic phytobiome is still a relatively unexplored realm, and there are several technical barriers preventing its widespread exploration. Sequencing studies can be complicated by the vast differences in 18S rRNA gene copy number among protists (Gong and Marchetti 2019), or the fact that even the most “universal” of primers introduce bias and miss some taxa (Adl et al. 2013). Functional studies can be hampered by the difficulty in isolating and culturing many soil eukaryotes. In this study we developed a tool to overcome one important hurdle in profiling the eukaryotic phytobiome, and used a pipeline that others can implement to develop their own similar tools. We hope this will encourage the phytobiome research community to incorporate a broader spectrum of rhizosphere and phyllosphere organisms into routine studies, as well as into discussions about what standardized sequencing methods to adopt.

## Acknowledgements

This work was supported by an AFRI Foundational Program grant from the USDA National Institute of Food and Agriculture to LRT, DJG, and BS (grant no. 2019-67019-29315). Kate Manning and Carlos Calderon provided technical and field assistance, and were supported through an AFRI Education and Learning Initiative grant from USDA-NIFA (grant no. 2017-67032-26013). We thank Drs. Jan Leach and Courtney Jahn, and Stephen Dellaporta and Ms. Emily Luna for contributing seeds of maize, wheat, barley and sorghum.

## List of Supplemental Files

Fig. S1. Histograms of matches between maize-derived 15, 16 and 17 bp k-mers and sequences from the corresponding V4 and V9 regions in a SILVA eukaryotic database that excluded plant sequences. Red arrows indicate the position of the selected PNA clamps (PoacV4_01 and PoacV9_01). X-axis units represent individual k-mers. Y-axis values represent the number of matching SILVA sequences, cut off at 100.

Fig. S2. Agarose gel image showing amplification of plant and protist V4 and V9 regions with and without addition of 1.5 mM of the corresponding PNA clamp.

Table S1. K-mers that passed screening as candidate clamps for suppression of maize DNA amplification. K-mer sequence, orientation, and the number of plant species with a matching target sequence are listed.

Table S2. List of plant species that are predicted *in silico* to be blocked by PoacV4_01 and PoacV9_01.

Table S3. Mean cycle thresholds (number of PCR cycles until detectable fluorescence) for amplification of the maize V9 region with different concentrations of clamp PoacV9_01 added. The means are shown with their ranges in parentheses.

Table S4. Relative abundances of the major taxonomic groups in clamp and no-clamp libraries, as classified by the PR^2^ and SILVA databases. Values are expressed as the proportion of total reads.

Table S5. Relative abundances of the main taxa within the fungi, animals and protists in clamp and no-clamp libraries, as classified by the PR^2^ databases. Values are expressed as the proportion of total reads.

Table S6. Read numbers of animals, fungi and protists in clamp and no clamp libraries as classified by the PR^2^ database.

## Notes

### Competing Interest Statement

The authors have declared no competing interest.

