## Supplementary figures and images for "Validation of a PNA clamping method for reducing host DNA amplification and increasing eukaryotic diversity in rhizosphere microbiome studies"

### Fig. S1

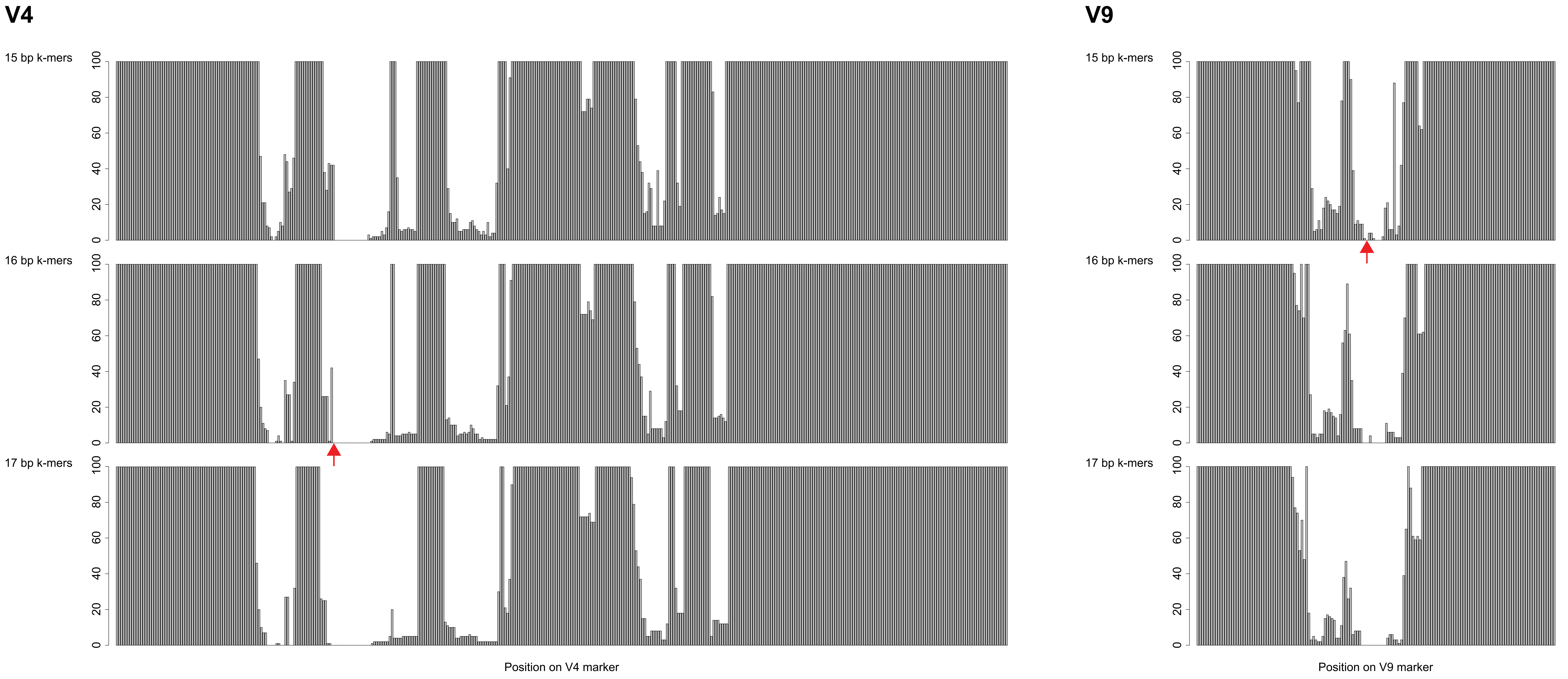
