## Supplementary material for "Validation of a PNA clamping method for reducing host DNA amplification and increasing eukaryotic diversity in rhizosphere microbiome studies": Fig. S2

### Slide 1
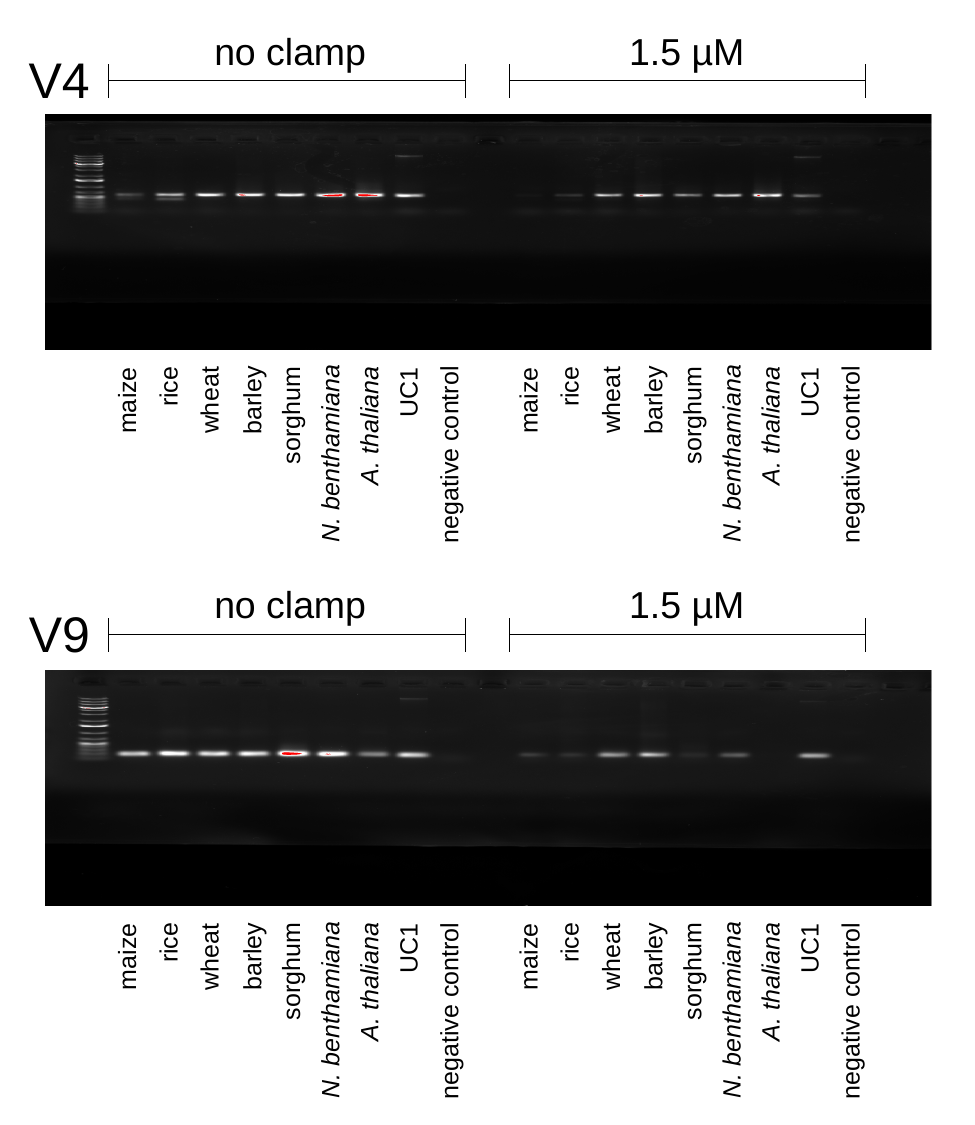

no clamp
1.5 µM
V4
rice
rice
UC1
UC1
maize
wheat
barley
maize
wheat
barley
sorghum
sorghum
A. thaliana
A. thaliana
N. benthamiana
N. benthamiana
negative control
negative control
no clamp
1.5 µM
V9
rice
rice
UC1
UC1
maize
wheat
barley
maize
wheat
barley
sorghum
sorghum
A. thaliana
A. thaliana
N. benthamiana
N. benthamiana
negative control
negative control
